# Post-proline cleaving enzymes also show specificity to reduced cysteine

**DOI:** 10.1101/2024.07.12.603020

**Authors:** Zuzana Kalaninová, Jasmína Mária Portašiková, Barbora Jirečková, Marek Polák, Jana Nováková, Daniel Kavan, Petr Novák, Petr Man

## Abstract

In proteomics, post-proline cleaving enzymes (PPCEs) like *Aspergillus niger* prolyl endopeptidase and neprosin complement proteolytic tools because proline is a stop site for many proteases. However, our systematic analysis of cleavage preferences showed that both PPCEs also display specificity to reduced cysteine. Post-cysteine cleavage was blocked by Cys alkylation, explaining why this activity has remained undetected. Our findings redefine their applicability and how we study and interpret their cleavage mechanism.

---

Proteolysis is a key step in classical bottom-up and structural proteomics and mass spectrometry-based structural analysis of proteins by H/D exchange (HDX), chemical cross-linking and radical labeling^1–3^. Protein digestion into shorter peptides facilitates mass spectrometric analysis and data processing, so this approach is much more widespread than intact protein analysis. Most often, these studies rely on specific enzymes with a predictable digestion pattern, particularly trypsin.

Trypsinization has long been the gold standard of proteolysis because trypsin cleaves after Lys and Arg with minimal missed sites. Yet, despite its reliability and numerous other advantages^4^, unwavering research interest in alternative enzymes for proteomics^1^ has provided access to proteins with an unfavorable Lys and Arg distribution^1,4–6^ and with post-translational modifications^7,8^. Among these promising enzymes, two post-proline cleaving enzymes (PPCEs) stand out for their highly selective cleavage after proline and alanine^9^, namely a glutamate endopeptidase termed neprosin^6^ and a member of the serine peptidase family S28^10^, *Aspergillus niger* prolyl endopeptidase (*An*PEP)^11,12^.

*An*PEP specificity can be fine-tuned by tightly controlling digestion conditions^5^. So, while aiming at using *An*PEP in online proteolysis, we found that this enzyme also displayed specificity to reduced cysteine. Through LC–MS/MS analysis, we confirmed this previously overlooked cleavage preference by systematically analyzing both *An*PEP sources and conditions that could affect this preference. And in the same experimental paradigm, neprosin mimicked this cleavage specificity. Based on these results, PPCEs cleavage preferences should be redefined from post-Pro/Ala to post-Pro/Ala/Cys. Moreover, this evidence demands reconsidering PPCEs applications, whether cleaving Cys-rich proteins or assessing Cys status in proteins, and calls for revisiting the proposed enzymatic mechanism of these proteases^13–15^.

Following recent reports describing *An*PEP/ProAlanase^5^ as a specific enzyme, we prepared *An*PEP columns for online protein digestion using an inexpensive Clarity Ferm-based material as it showed slightly better quality than the Tolerase G and was easier to handle (Extended Data Fig. 1). We tested the column in a typical hydrogen/deuterium exchange LC-MS setup, replacing the digestion/desalting solution by 30mM hydrochloric acid to lower the pH to 1.5, which is crucial for *An*PEP specificity^5^. A mixture of pre-reduced purified proteins was digested online under different flow rates (100 and 200 μL/min) and temperatures (2 and 21 °C), thus modulating the digestion conditions^16^.

By analyzing cleavage preferences, we found that the immobilized enzyme was not as specific as expected (Extended Data Fig. 2). Only when considering fully reproducible peptides (6 of 6) were the preferences more centered around the expected post-Ala and Pro specificity (Fig. 1a). More stringent conditions (lower temperature and faster flow) had a similar effect, but other cleavage sites (post-Arg, Cys, Ser, Lys and Gly) were also significant. For this reason, we digested reduced human serum to test these observations in a more complex setting.

**Fig. 1.**
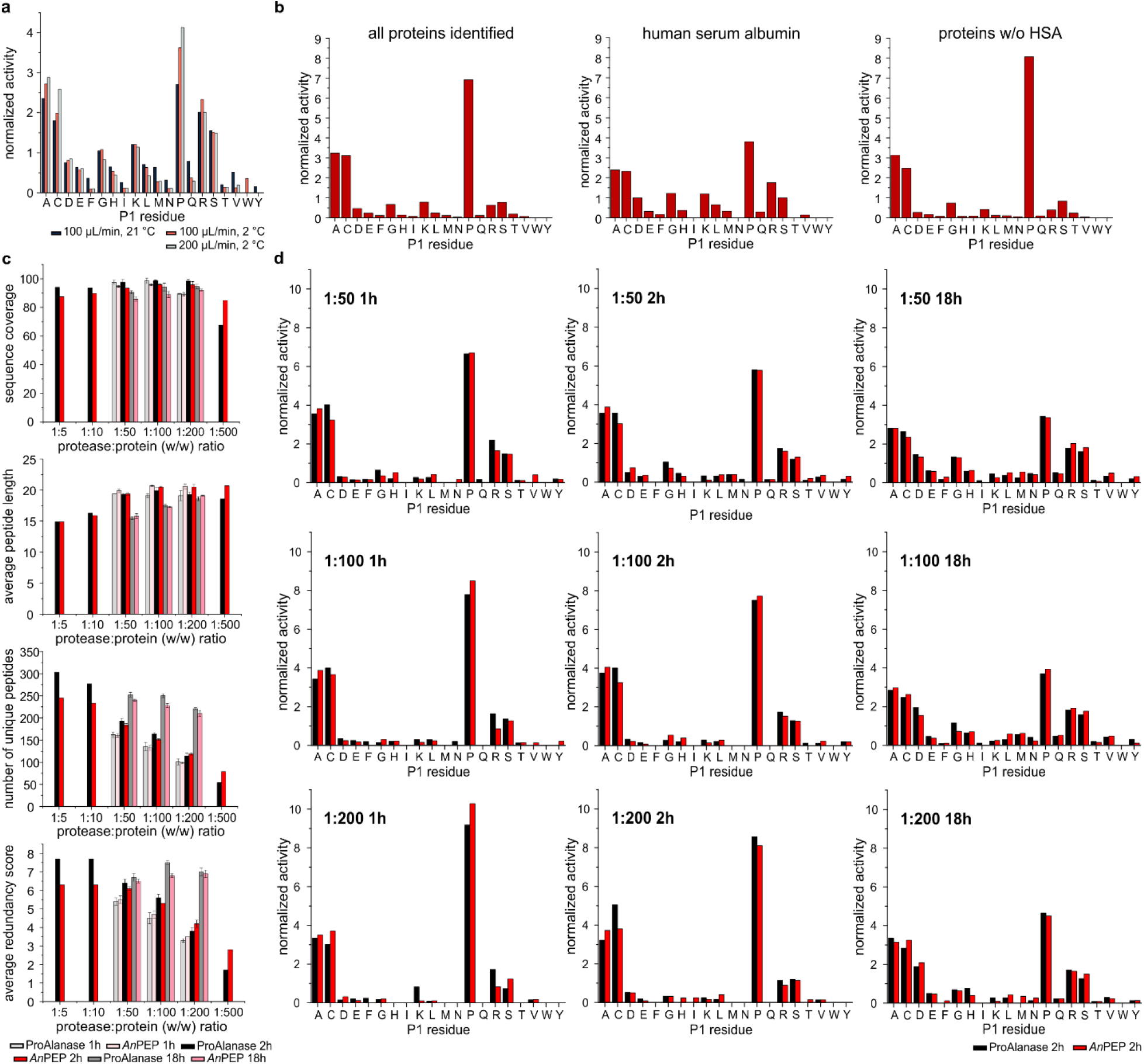
ProAlanase and *An*PEP cleavage preferences and digestion parameters match and highlight post-Cys cleavage activity. **(a)** Cleavage preferences for online digestion of a protein mixture on an *An*PEP-immobilized column at various temperatures and flow rates. **(b)** Online digestion of serum on an *An*PEP-immobilized column, differences in cleavage preferences of all identified proteins, top hit human serum albumin and all identified proteins without HSA. **(c)** Key digestion parameters (sequence coverage, average peptide length, number of unique peptides and average redundancy score) and **(d)** cleavage preferences of ProAlanase (black) and *An*PEP (red) in solution digestions at different time points and enzyme:protein ratios.

Preferences built on fully reproducible peptides showed a much higher specificity, with post-Pro/Ala and, surprisingly, post-Cys being, by far, the most common cleavage sites. For serum albumin, the most abundant plasma protein, the extracted preferences were less selective, matching those observed in a test mixture. Nevertheless, the post-Pro/Ala/Cys prevalence held for the remaining serum proteins (Fig. 1b). Therefore, we should not rely on only partial sequence coverages, limited to the most prevalently generated peptides, for data processing; otherwise, we may overestimate cleavage specificity (Fig. 1b – preferences without HSA). Case in point is the high preference of *An*PEP for cleavage after Cys residues.^5,11,14^

To ascertain whether the lower selectivity and Cys-directed preference of immobilized *An*PEP could be attributed to immobilization or were inherent properties of the Clarity Ferm-based material, we compared *An*PEP and ProAlanase (MS-grade *An*PEP) digestions in solution. For this purpose, we monitored the digestion of a pre-reduced protein mixture at 37 °C, pH 1.5, and enzyme:protein ratios of 1:50, 1:100, 1:200 for 1, 2, and 18h. In addition, we tested 1:5, 1:10, 1:500 ratios, but only for the optimal digestion time (2 h)^5^.

Built on fully reproducible peptides, the key digestion metrics of Clarity Ferm *An*PEP and ProAlanase showed a good agreement (Fig. 1c). Coverage was higher than 85% for the three main enzyme:protein ratios. Peptide count increased and length decreased with protease loading and incubation time. Cleavage preferences also matched, displaying logical trends (Fig. 1d and Extended Data Fig. 3 and 4).

Both enzymes were more specific at shorter digestion times and higher enzyme:protein ratios. In line with published data^5^, these two *An*PEPs predominantly showed post-Pro cleavage specificity, followed by a lower prevalence of fragments resulting from post-Ala cleavage. However, post-Cys cleavage was also significant in both enzyme preparations. Therefore, we ruled out immobilization effects or specific features of the Clarity Ferm material.

We also ruled out potential protein contaminations by proteomic profiling of Clarity Ferm *An*PEP. *An*PEP was the main component, corroborating the SDS-PAGE profile (Extended Data Fig. 1). Other minor components included alpha-amylase, glycosidases, and serine carboxypeptidase. But even serine carboxypeptidase could not be the source of minor, non-specific cleavages increasingly apparent at lower enzyme:protein ratios. Its activity can only account for ragged C-terminal ends^17^, not for peptides derived from an endopeptidase such as those in datasets with Cys preceding their N-terminal side. Accordingly, the lower specificity of the enzymatic columns may be attributed to high local protease concentration, so post-Cys activity is a novel specificity of *An*PEP.

This post-Cys activity of *An*PEP has not been detected previously^5,11,17^, most likely due to Cys alkylation. To test this hypothesis, we digested an alkylated protein mixture and compared the results with the corresponding data from the reduced protein mixture (Fig. 2a). The sharp decrease in post-Cys cleavage and the concomitant increase in post-Ala and post-Pro cleavages demonstrated that *An*PEP can only cleave the peptide bond C-terminal to reduced Cys and that Cys alkylation prevents this cleavage.

**Fig. 2.**
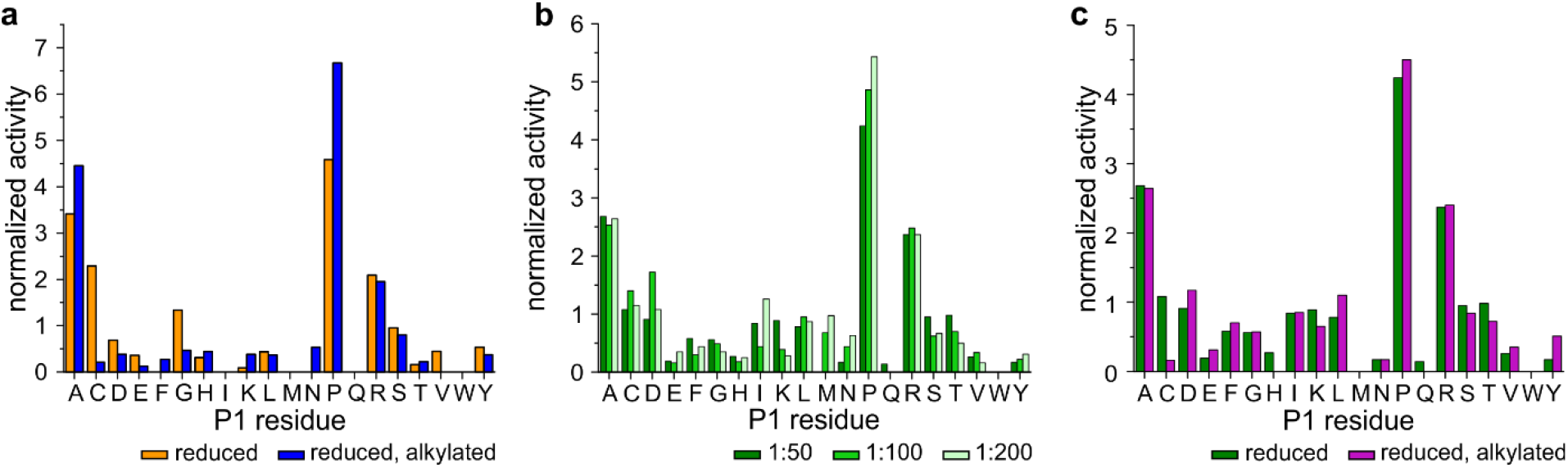
*An*PEP and neprosin cleavage preference for reduced cysteine is blocked by alkylation. **(a)** Differences in cleavage preferences of *An*PEP digestion of reduced (orange) and alkylated (blue) protein mixtures. **(b)** Cleavage preferences of neprosin followed at various enzyme:protein ratios. **(c)** Neprosin post-Cys cleavage is also blocked by alkylation, as shown by digestion of reduced (green) and alkylated (purple) protein mixtures.

We further tested *An*PEP digestion of a model disulfide-linked peptide – insulin – and its oxidized beta-chain, with and without Cys reduction. Non-reduced insulin was poorly digested, with post-Ala and Pro digestion yielding only minor fragments (Extended Data Fig. 5), as with oxidized beta-chain. In contrast, insulin digestion after Cys reduction was nearly complete and accounted for most post-Cys fragments. This evidence confirms that neither oxidized Cys nor Cys involved in disulfide bonds is cleaved by *An*PEP, supporting previous claims^5^.

Although neprosin is a glutamate endopeptidase and, as such, belongs to a different enzyme class from *An*PEP (prolyl endopeptidase), we found that its cleavage preferences can also be described as post-Pro<Ala<=Cys (Fig. 2b). Neprosin digestions were performed at pH 2.5, though, considering the pH optimum of this protease^6^, but as with *An*PEP, the post-Cys activity was also blocked by Cys alkylation (Fig. 2c). Once again, this Cys alkylation effect may explain why PPCE post-Cys activity has never been observed before^6^.

Overall, our results unveil the cleavage preference of *An*PEP and neprosin for reduced cysteine, mirroring the selectivity of human prolyl endopeptidase and dipeptidyl peptidases once suggested in a conference paper^18^. Because their post-Cyst cleavage has remained undetected in the proteomics field, we must reassess the impact of this activity on regulatory mechanisms in the human body^18^ and reconsider both PPCE applications^12,19^ and mechanisms of action^10,12^, thereby opening new opportunities in protein analysis. Since *An*PEP and neprosin digestion requires acidic conditions, proteins may be used in their reduced form, avoiding the alkylation step. These enzymes may also help to analyze the cysteine status of disulfide bond-containing proteins, such as antibodies^5^. Ultimately, studies defining enzyme preferences should use substrates representing all common amino acids in their natural, free form^20,21^.

## Methods

### Protease and sample preparation

Research grade ProAlanase was purchased from Promega. *Aspergillus niger* prolyl endoprotease (*An*PEP) was isolated from Clarity-Ferm (White Labs). A P-10 desalting column packed with Sephadex G-25 resin (GE Healthcare) was used for buffer exchange. The column was washed and equilibrated with water (4 × 4 mL) and sodium citrate buffer (50 mM, pH 5.0; 5 × 4 mL). Subsequently, 2.5 mL of Clarity-Ferm was applied and allowed to enter the bed completely. *An*PEP was eluted with 3.5 mL of sodium citrate buffer. Protein concentration was determined using a Pierce BCA protein assay kit (Thermo Fisher Scientific). Isolated *An*PEP was immobilized on an aldehyde-functionalized POROS 20 AL resin, as described previously.^22–24^ Recombinant neprosin was expressed according to a literature protocol.^6^

A protein mixture in Tris-HCl buffer (100 mM, pH 8.8) was used for offline and online digestion experiments. This mixture consisted of five proteins, namely bovine serum albumin (BSA), bovine carbonic anhydrase II (bCA2), horse cytochrome C (cytC) and horse myoglobin (Mb), all of which purchased from Merck Life Sciences, and 14-3-3gamma, which was expressed recombinantly.^25^ The concentration of all proteins was 10 μM, as confirmed using the Pierce BCA protein assay kit. The mixture was reduced with either 10 mM tris-(2-carboxyethyl)-phosphine (TCEP) at 65 °C for 10 min. The Cys alkylated sample was prepared by incubating the reduced proteins with 20 mM iodoacetamide (IAA) for 1 h at room temperature in the dark. Human insulin and oxidized bovine beta-insulin chain were purchased from Merck Life Sciences. Serum from healthy donors of the Department of transfusion medicine and blood bank at Thomayer University Hospital in Prague was diluted (10 ×) with Tris-HCl buffer (100 mM, pH 8.8) and reduced with TCEP (10 mM) for 10 min at 65 °C.

### Insulin digestion and MALDI-FT-ICR MS analysis

Insulin was dissolved in 5% acetic acid, and its disulfide bonds were reduced using 50mM TCEP for 4 h at 21 °C. The oxidized beta chain was dissolved directly in 50mM Tris-Cl pH 8.0. Non-reduced and reduced insulin and oxidized beta chain were transferred to 30mM HCl pH 1.5, followed by *An*PEP digestion at a 1:50 enzyme:protein ratio for 18 h at 37 °C. Samples were analyzed on a Matrix-assisted laser desorption/ionization Fourier transform ion cyclotron resonance mass spectrometry (MALDI-FT-ICR MS; 15T solariXR, Bruker Daltonics) using alpha-cyano-4-hydroxycinnamic acid as the MALDI matrix.

### On-column protein digestion and LC-MS/MS analysis

Online digestion and analysis were performed in a fully automated mode. The samples were handled by a PAL DHR robot (CTC Analytics AG) controlled by the Chronos software (Axel Semrau). A 5 μL aliquot of reduced protein mixture (50 pmol per injection) was diluted with 95 μL of 30 mM HCl and manually injected into the LC system. To analyze serum samples, 10 μL of reduced serum was diluted with 40 μL of 50 mM HCl and mixed with 50 μL of 30 mM HCl or 4 M urea in 30 mM HCl. The LC system consisted of a custom-made column with immobilized *An*PEP (bed volume 66 μL), a trap column (SecurityGuard™ ULTRA Cartridge UHPLC Fully Porous Polar C18, 2.1 mm; Phenomenex), and an analytical column (Luna Omega Polar C18, 1.6 μm, 100 Å, 1.0 × 100 mm; Phenomenex). Sample digestion and peptide desalting were performed in 30 mM HCl delivered by the 1260 Infinity II Quaternary pump (Agilent Technologies) at 100 (2 or 21°C) or 200 (2°C) μL/min. The peptides were separated for 6 min in a water:acetonitrile gradient (5%-45% (v/v); solvent A: 0.1% FA in water, solvent B: 0.1% FA, 2% water in acetonitrile) followed by a step to 99% B delivered by the 1290 Infinity II LC system (Agilent Technologies) at a flow rate of 40 μL/min. Serum samples were analyzed for 27 min in a water:acetonitrile gradient (5%-35% (v/v); solvent A: 0.1% FA in water, solvent B: 0.1% FA, 2% water in acetonitrile) followed by a step to 99% delivered by the 1290 Infinity II LC system. The timsTOF Pro mass spectrometer with active parallel accumulation–serial fragmentation (PASEF) and 100 ms tims ramp time was operated in a data-dependent MS/MS mode.

### In-solution protein digestion and analysis

To test protease activity in-solution, 5 μL aliquots of reduced protein mixture (containing 15 μg of protein) were diluted with 95 μL of HCl (30 mM, pH 1.5). Protease (1 μL), either research grade ProAlanase or purified *An*PEP, was added at a 1:50, 1:100 and 1:200 ratio (w/w). The reactions were performed in triplicate for 1, 2, and 18 h digestion. Additionally, reaction mixtures at 1:5, 1:10 and 1:500 (w/w) enzyme:protein ratios were incubated for 2 h only. An alkylated protein mixture was used to investigate cleavage preferences related to cysteine status by digestion for 2 h at a 1:50 (w/w) ratio in triplicate. All digestions were stopped by increasing the pH to 7.0 with 1 M Tris solution.

Samples were subsequently analyzed by LC-MS/MS. Peptides were first desalted by 0.4% formic acid (FA) in water driven by an LC-20AD pump (Shimadzu) at 20 μL/min using a Luna Omega (5 μm Polar C18 100 Å, Micro Trap 20 × 0.30 mm) trap column (Phenomenex) and then eluted and separated for 15 min in a water:acetonitrile gradient (5%-35% (v/v); solvent A: 0.1% FA in water, solvent B: 0.1% FA, 2% water in acetonitrile), followed by a step to 99% solvent B driven by the Agilent 1200 HPLC system at a flow rate of 4 μL/min using a Luna Omega (3 μm Polar C18 100 Å, LC Column 150 × 0.30 mm) analytical column (Phenomenex) heated to 50 °C. The LC system was directly connected to an ESI source of the timsTOF Pro mass spectrometer with enabled PASEF (Bruker Daltonics), operating in a data-dependent MS/MS mode with a 100 ms tims ramp time.

To test neprosin, another post-Pro/Ala protease, for its preference for Cys, recombinant protease was added to 5 μL of protein mixture diluted with glycine (50 mM, pH 2.5) at 1:50, 1:100 and 1:200 (w/w) ratios, and the reactions were performed for 2 h at 37 °C in triplicate. The pH was raised to 7.0 with 1 M Tris, the peptides were desalted, eluted and separated for 15 min in a water:acetonitrile gradient (4%-35% (v/v); solvent A: 0.1% FA in water, solvent B: 0.1% FA, 20% water in acetonitrile), followed by a step to 99% solvent B driven by the UHPLC system (VanquishTM Neo, Thermo Scientific) at a 1.5 μL/min flow rate. The system consisted of a PepMap Neo (5 μm C18, 50 × 0.30 mm, Thermo Scientific) trap cartridge and a PepSep (1.5 μm C18, 150 × 0.15 mm) analytical column heated to 50 °C. The outlet was directly coupled with an ESI source of a timsTOF SCP mass spectrometer (Bruker Daltonics), operating in a data-dependent mode employing PASEF.

### AnPEP profiling by LC-MS/MS

Clarity Ferm *An*PEP was deglycosylated by Endo H (50 mM Bis-Tris-Cl pH 6.0, 37 °C, 18 h), and subsequently transferred into 50 mM ammonium bicarbonate and subjected to in-solution digestion by either trypsin or Glu-C or AspN. Cys residues were blocked by reduction with 10 mM TCEP and by alkylation with 20 mM iodoacetamide. The peptides were injected onto a PepMap Neo (5 μm C18, 50 × 0.30 mm, Thermo Scientific) trap cartridge, where they were desalted with 0.1% formic acid in water and separated on a PepSep (1.5 μm C18, 150 × 0.15 mm) analytical column for 15 min by a water:acetonitrile gradient (4%-35% (v/v); solvent A: 0.1% formic acid in water, solvent B: 0.1% formic acid, 20% water in acetonitrile). The column was kept at 50 °C. Solvents were delivered by the UHPLC system (VanquishTM Neo, Thermo Scientific) at a 1.5 μL/min flow rate. The outlet of the LC was directly coupled to an ESI source of timsTOF SCP mass spectrometer (Bruker Daltonics), operating in a data-dependent mode employing PASEF.

### Data processing

LC-MS/MS data were peak picked in DataAnalysis 5.3 software (Bruker Daltonics), exported to *.mgf files and searched using MASCOT (version 2.7, Matrix Science) against a custom-built database combining the sequences of the proteins in the mixture with a common cRAP.fasta (https://www.thegpm.org/crap/) and the sequences of the proteases. Serum digestion data were searched against the UniProt/Swiss-Prot database (release 2023_01). *An*PEP profiling data were searched against a subset of trEMBL database containing all proteins from *Aspergillus sp*. and a specific sequence of *An*PEP from GenBank (GAQ41715.1). In all cases, the MASCOT search parameters were set to 10 ppm precursor tolerance, 0.05 Da fragment ion tolerance and decoy search enabled, with FDR < 1%, IonScore > 25, and peptide length > 6. Variable modifications: Heme or Dehydro (Cys), acetylation (protein N-term) were used for protein mixture. Carbamidomethylation (Cys) was added for alkylated samples. *An*PEP profiling data included carbamidomethylation (Cys) as a fixed modification, while acetylation (protein N-term) and HexNAc (Asn) were entered as variable modifications. Full MASCOT search results were exported as *.csv files. Digest parameters – average redundancy, redundancy distribution, sequence coverage, number of unique peptides, average peptide length, and cleavage preferences – were evaluated using a new in-house developed Java-based software tool, DigDig (can be obtained from authors upon request).

**Supplementary data** are provided as a separate document associated with this manuscript. All primary MS data and secondary data (*.mgf, *.csv, databases) have been deposited to ZENODO under DOI: 10.5281/zenodo.12655607

## Acknowledgments

We thank Petr Turek, MD, Csc, Department of Transfusion Medicine and Blood Bank, Thomayer University Hospital, for the serum samples. This study was funded by the National Institute for Neurological Research (Programme EXCELES, ID Project No. LX22NPO5107), supported by the European Union – Next Generation EU, and by the Technology Agency of the Czech Republic (Programme National Centre of Competence, ID Project No TN02000132 National Centre for new methods of diagnosis, monitoring, treatment and prevention of genetic diseases). Access to the Centre of Molecular Structure (CMS), BioCeV, was provided by CIISB LM2023042 and ERDF “UP CIISB” (CZ.02.1.01/0.0/0.0/18_046/0015974). We thank Carlos V. Melo for helpful discussions and manuscript editing.

## Author contributions

Z.K. performed the experiments, analyzed the MS data, prepared the figures and contributed to the drafting of the manuscript, J.M.P. and B.J. performed the initial experiments, M.P. and J.N. prepared the immobilized protease columns and contributed to pilot studies, D.K. wrote scripts for data processing and carried out bioinformatic work, P.M. designed the study, supervised the project, validated data processing, prepared the manuscript, P.N. provided grant support and contributed to manuscript preparation and project supervision. All authors edited and approved the final manuscript.

## Competing financial interests

Petr Novak is one of the founders of the AffiPro s.r.o.

