## Supplementary material for "Post-proline cleaving enzymes also show specificity to reduced cysteine"

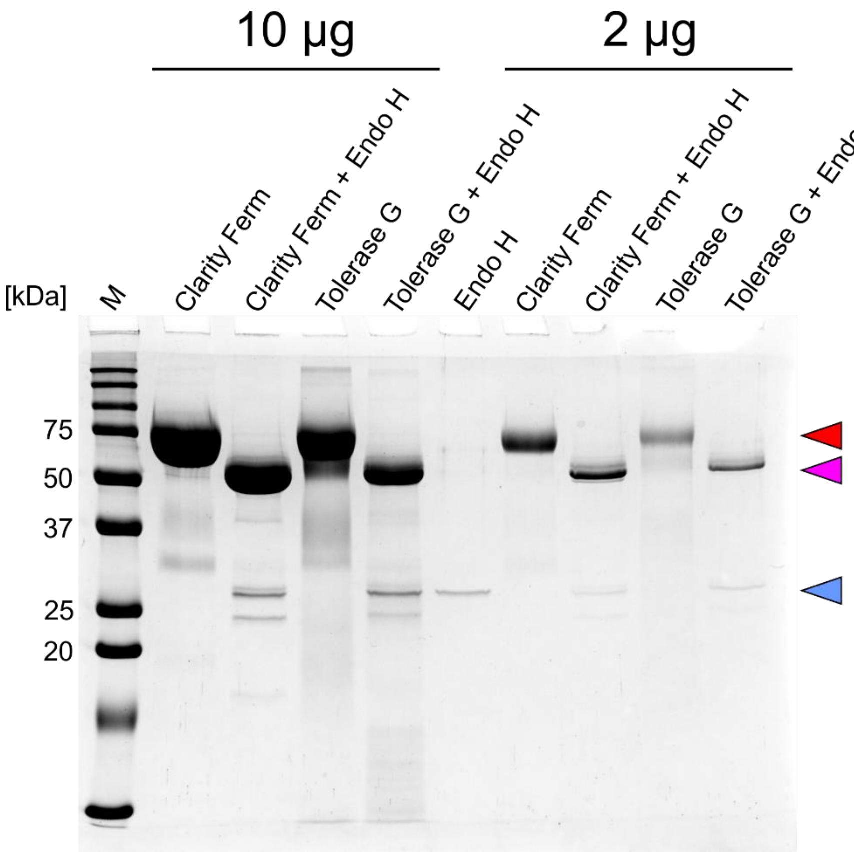


**Extended Data Fig. 1.** SDS-PAGE analysis of two different *An*PEP sources – Clarity Ferm and Tolerase G – before (red triangle) and after (pink triangle) deglycosylation with EndoH (blue triangle). Both sources show high purity/homogeneity, with only minor contaminations and are highly N-glycosylated. Two protein amounts, 10 and 2µg, were used, as indicated above the gel.

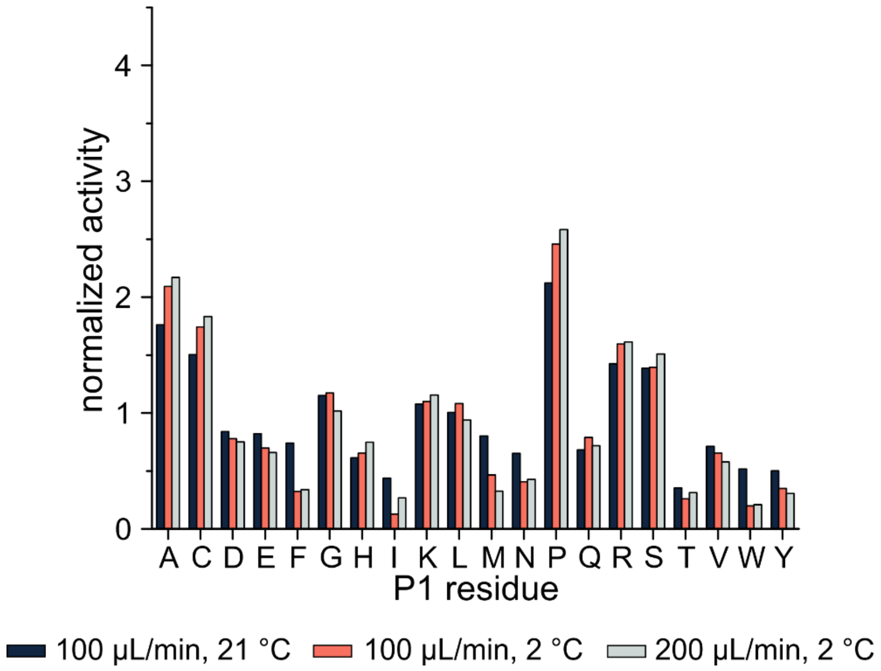


**Extended Data Fig. 2.** Cleavage preferences for online digestion of a protein mixture on an *An*PEP column, considering all identified (not just the fully reproducible) peptides.

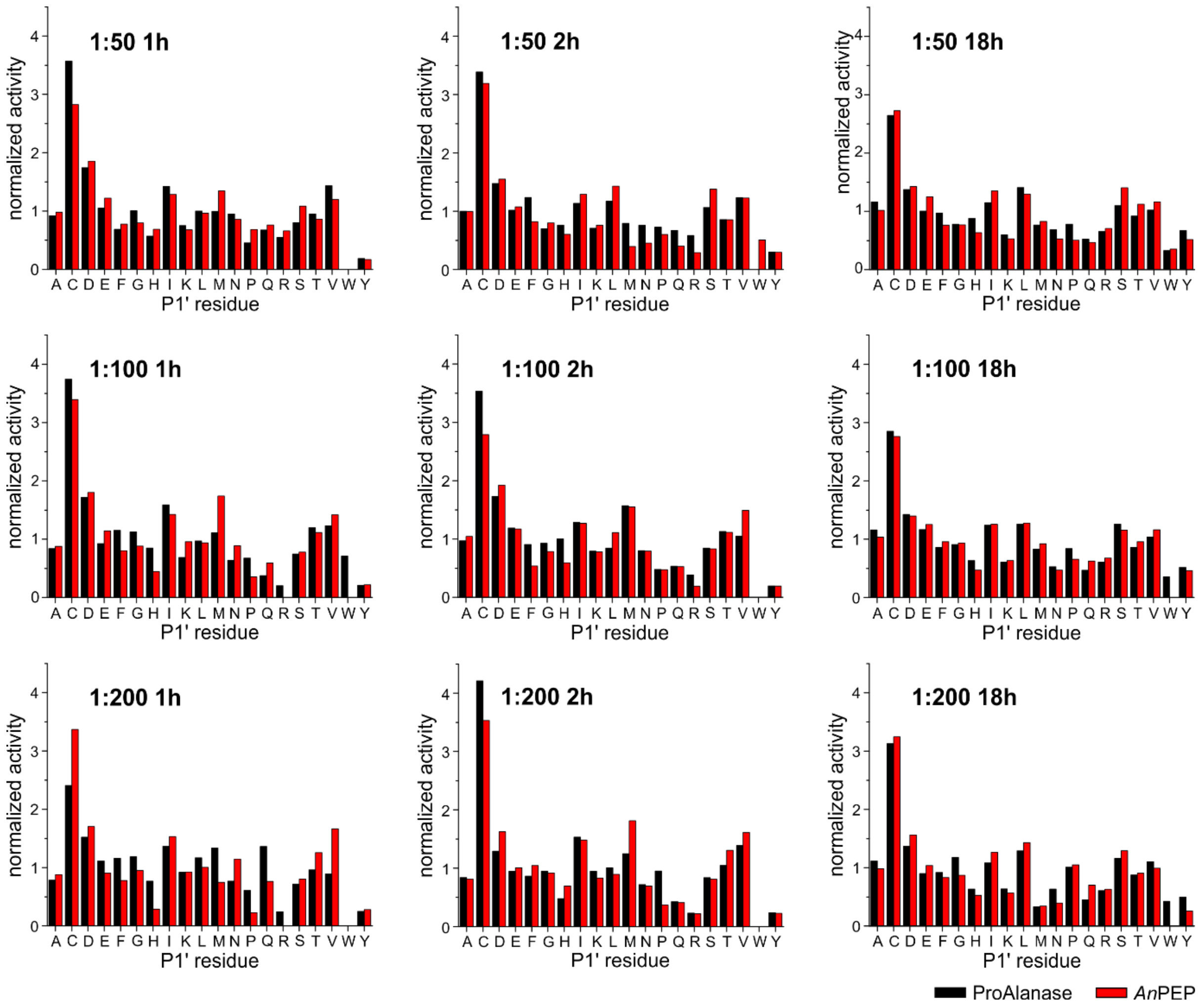


**Extended Data Fig. 3.** Cleavage preferences at the P1′ site for in-solution digestion of a protein mixture comparing *An*PEP and ProAlanase at different digestion times (1, 2 and 18 h) and enzyme:protein ratios (1:50, 1:100 and 1:200). All graphs are normalized to the same value at the y-axis.

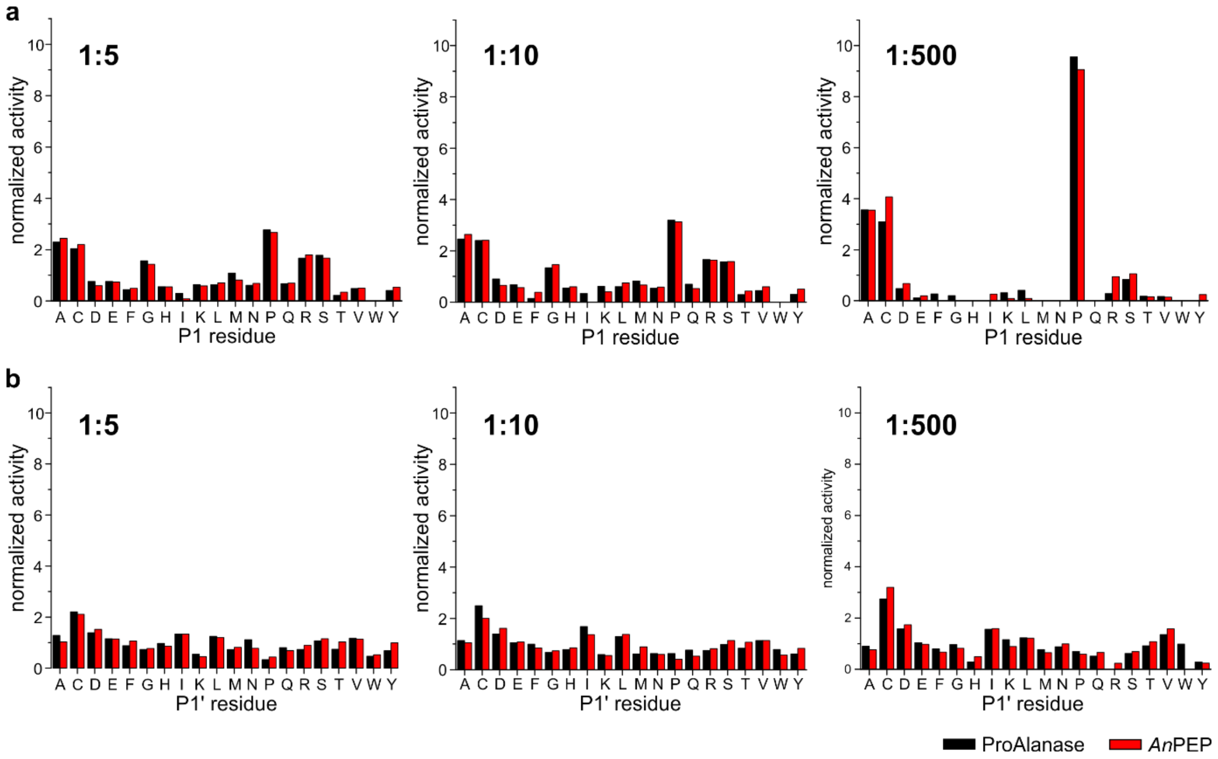


**Extended Data Fig. 4**. Cleavage preferences at P1 (**a.**) and P1’ (**b.**) site for in-solution digestion of a protein mixture comparing *An*PEP and ProAlanase at 2-hour-digestion time and 1:5, 1:10 and 1:500 enzyme:protein ratios. All graphs are normalized to the same value at the y-axis.

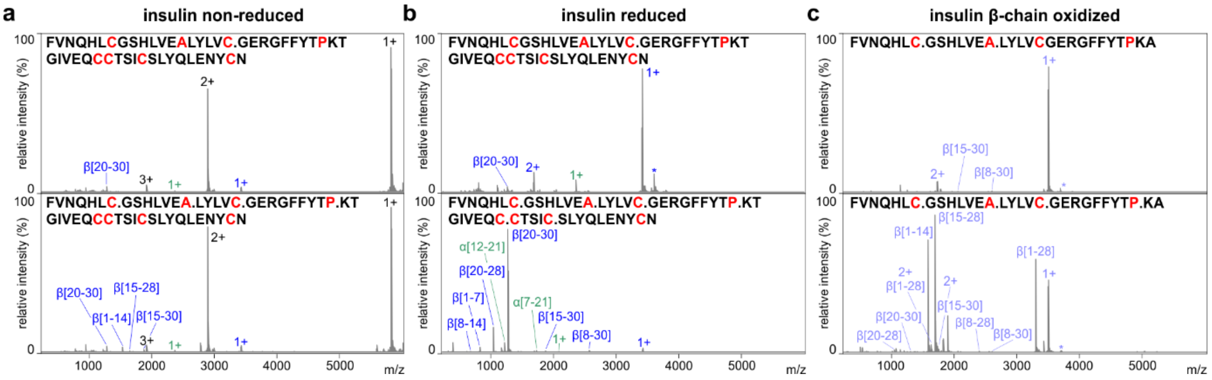


**Extended Data Fig. 5.** *An*PEP digestion of **(a)** non-reduced and **(b)** reduced human insulin and **(c)** oxidized bovine insulin beta-chain. Spectra before (top) and after (bottom) AnPEP digestion for 2 h at 37 °C and a 1:50 enzyme:protein ratio. Intact insulin is highlighted in black; fragments from the alpha chain, in green; and fragments from the beta chain, in blue.
